# Temporal characterization of the viral load of psittacine beak and feather disease virus in rosy-faced lovebirds (*Agapornis roseicollis*)

**DOI:** 10.1101/2024.02.19.581011

**Authors:** Derek Kong Lam, Emily Shui Kei Poon, Simon Yung Wa Sin

## Abstract

Psittacine beak and feather disease virus (PBFDV) is a widespread and highly pathogenic virus in parrots (Psittaciformes), threatening both captive and wild populations over the world. The disease typically presents with feather and beak abnormalities, along with possible immune system suppression. No cure or commercialized vaccine is currently available. Our understanding of the Psittacine beak and feather disease often come from infected individuals with visible symptoms. Limited knowledge exists regarding the pathology and role of asymptomatic individuals in disease transmission. Asymptomatic individuals could shed virus in their crop secretion, feces, or feathers. In this study, we investigated the temporal change in viral load in feather and fecal samples from 17 asymptomatic rosy-faced lovebirds (*Agapornis roseicollis*). We developed a qPCR assay for PBFDV viral load quantification in the studied lovebirds. Our results showed that most of the individuals had very low viral load, while three individuals with high viral load at the beginning of the experiment were observed to exhibit a decreasing trend in viral load in both fecal and feather samples. Surprisingly, the viral load in an individual can drop from a high level to an undetectable level within three months, which is contrary to the prevailing notion that the disease is highly lethal with few reports of complete recovery. We also showed that viral load in feathers was higher than in feces. Our study provides valuable insights into the infection dynamics of PBFDV in asymptomatic individuals and contribute to the understanding of disease transmission in parrots.

## Introduction

Psittacine beak and feather disease (PBFD) poses a major threat to both wild and captive parrots worldwide [8, 38, 39]. The disease is caused by the highly contagious psittacine beak and feather disease virus (PBFDV), a single-stranded DNA circovirus [2, 40]. The virus can infect most, if not all, psittacine species, which is particularly concerning due to the endangered status of many parrot species [8, 10, 21, 44, 51]. The global trade of parrots has been shown to play a significant role in disseminating the virus across different continents, including Asia, Africa, the Americas, Europe, and Australasia [3, 4, 9, 11, 12, 26]. Recent reports also highlighted the increasing PBFDV occurrence in non-psittacine species [1, 36]. Although the pathogenicity of PBFDV in non-psittacine species remained to be determined, it is also concerning that those infected non-psittacine species could play a role in disease transmission.

The infection outcomes of PBFD in parrots can be severe and lethal. Acute cases were frequently observed in young individuals with a sudden death without visible symptoms [48, 19]. Chronic PBFD tends to occur in older individuals with or without manifestation of symptoms. Once infected, the virus will remain in the host for the rest of its life, thus the infected individual could become carriers and may shed virus at any point of its life. Asymptomatic individuals may start developing visible symptoms once their immune system is weakened [40]. Typical symptoms of PBFD include feather dystrophy and abnormal beak and claw growth [31]. Besides, the virus is immunosuppressing, targeting the immune system, particularly the thymus and bursa of Fabricius, thus leading to depletion of lymphocytes [48]. Infected individuals often die from secondary infections [48].

To date, despite the discovery of PBFD in Australia in the 1970s, no cure or effective treatment has been discovered for the disease [32]. Vaccines for PBFDV has been recently developed but not yet commercialized. In captive environment, the major measures for controlling PBFDV infection are better hygiene and early diagnosis, usually performed using polymerase chain reaction [PCR; 10, 11]. Infected individuals, even asymptomatic, could shed virus in their crop secretion, feces, or feathers, which contaminate the environment [41]. Transmission mechanism of the PBFD include direct contact, fecal-oral route, vertical transmission (i.e., from the parent to the offspring), contaminated water and food, feather and skin particles [22,37]. Therefore, infected individuals, even asymptomatic, need to be isolated from healthy individuals permanently, which might pose a burden for captive population management especially for breeding industry.

Over the three decades, the focus of PBFD research has been shifted from disease characterization to PBFDV screening, prevalence, evolution, and phylogenetic analysis [8, 14, 17, 18, 35]. A wealth of data on the PBFDV genotypes and prevalence are available [12, 13, 24, 28, 44, 49]. However, our understanding of the infection dynamic, which is critical for designing effective conservation management strategies and clinical treatments, is still limited. For example, numerous studies that use PCR detection have reported a high PBFDV prevalence in wild and captive parrot populations with most individuals being asymptomatic [1, 28, 33, 36]. However, the viral load and shedding patterns in these asymptomatic birds, as well as their relationships with the overall health conditions of the birds, are unclear [1]. Such knowledge could help estimating transmission rate, identifying high-risk individuals or populations for more targeted actions, and advance our understanding of the infection dynamic between the virus and the host. Furthermore, our knowledge on PBFDV pathology primarily came from acute individuals with autopsy examination [42, 45, 47]. The characteristics of the physiological changes (e.g., anemia, leukopenia, liver necrosis, etc.) in different parrot species has been reported but in acute individuals [45]. These pathology studies were limited by a biased sample size, as most of the samples used in previous studies came from clinics where the birds were severely ill. The impacts of the virus on asymptomatic and carrier individuals were completely unknown.

International trades, both legal and illegal, has been identified as the major factor contributing to the fast spread of PBFDV [49]. Among all the parrots, lovebirds (*Agapornis*) were the most traded genus according to the record of CITES, with the trade volume of 4,287,540 individuals between 1975 to 2016 [6]. The actual trade volume was even higher because the rosy-faced lovebird (*Agapornis roseicollis*) was removed from the CITES Appendices in 2005. *Agapornis* are small African parrots with nine existing species in the genus [16, 25]. Some of them are popular as companion pets, especially *A. roseicollis* [27]. Invasive populations of lovebirds have been reported in the UK, France, Italy, and Spain [27]. The outcomes of PBFDV infection in lovebirds ranged from sub-clinical to lethal. Some veterinarians suggested that most of the infected lovebirds in US are asymptomatic carriers. Given the high international trade volume, establishment of invasive population worldwide, and the popularity as a companion pet, lovebirds could serve a major reservoir of the disease [20, 50].

To this end, given very limited information on the role of asymptomatic individuals in PBFDV transmission, we aimed to investigate the temporal change in viral load in feather and fecal samples from 17 asymptomatic *A. roseicollis*, as feather and feces are the primary means of viral shedding. To achieve that, we developed a quantitative PCR (qPCR) assay for PBFDV DNA quantification in our infected birds since PBFDV genome is very variable with some studies showing host-specific viral sequences [5, 43]. Our findings provide valuable insights into the dynamics of host-virus infections and offer crucial information for disease and conservation management.

## Materials and methods

### Sample collection

Seventeen rosy-faced lovebirds (14 males and 3 females) obtained from local breeders were housed in the Centre for Comparative Medicine Research (CCMR) in the University of Hong Kong, and kept individually in individually ventilated cages (IVC; Tecniplast, Italy) to avoid cross-infection or contamination. The age at which the birds arrived at the facility varied between 2- to 23-month old. Around 7 feathers were plucked from each of following body regions: back, chest, and down feathers on the rump, using sterilized forceps with clean nitrile gloves. Fresh fecal samples were collected from the bottom of the IVC using forceps. In total, 66 fecal samples and 138 feather samples were collected. Samples were collected from each bird once per month on average. All samples were stored in 1.5-ml microcentrifuge tube at -80°C until DNA extraction. All samples were collected inside a biosafety cabinet.

### DNA extraction

DNA from feather samples were extracted using DNeasy Blood & Tissue kits (Qiagen, Germany) inside a biosafety cabinet. Both enzymatic and mechanistic treatments were included in the extraction processes. Samples were incubated in 360 µl enzymatic buffer (20 mM Tris-HCL, 2 mM sodium EDTA, 1.2% Trition X-100, 20 mg/ml lysozyme, at pH 8.0) at 37°C for 1.5 h with 8 rpm rotations. Two-hundred mg glass beads were then added into the samples. Samples were shaken vertically for 5 min at 30 Hz in a TissueLyser (Qiagen), followed by proteinase K treatment (50 µl) at 56°C for 30 min. Samples were then incubated at 95°C for 5 min. Next, samples were centrifuged to remove pellets and the supernatant was pipetted for DNA purification using DNeasy Mini spin column according to the manufacturer’s protocol. DNA from fecal samples were extracted using QIAamp Fast DNA Stool Mini Kit (Qiagen) inside a biosafety cabinet following the manufacturer’s protocol. All extracted DNA were stored at -80°C until further analysis.

### Quantitative PCR (qPCR)

Four hundred and twenty PBFDV sequences were downloaded from GenBank and aligned using MEGA7, together with another 14 PBFDV sequences isolated from our samples. Since the PBFDV sequences were highly variable, we focused on the PBFDV sequences isolated from our lovebirds (i.e. 20 sequences) for primer design. A pair of primers were designed on the relatively conserved region of the *rep* genes, which is one of the two protein-coding genes in the viral genome [30]. qPCR was performed in duplicate per sample in a 15-μl reaction mix containing 2X iTaq Universal SYBR Green Supermix (Bio-Rad, US), 0.4 μM forward primer (5’-GAGCTGTTGCTGCCGTGAT-3’), 0.4 μM reverse primer (5’-CGCCCATGCCTGACGTAG-3’), and 20ng gDNA using CFX96 Torch Real-Time PCR Detection System (Bio-Rad, US). The thermal cycle program consisted of 1 cycle of 2 min at 95°C, and 40 cycles of 5 sec at 95°C and 10 sec at 62°C, ended with a melt curve analysis ramping from 65°C to 95°C with 0.5°C/5 sec increment. We used standard curve method to calculate the absolute DNA copy number present in the sample. To construct the standard curve, we first cloned the amplicon into the T-vector pMD19 (TaKaRa Bio, Japan) using DNA ligation kit (TaKaRa). The resulting plasmids containing the PBFDV sequence were linearized using a restriction enzyme ScaI-HF (New England Biolabs, US) and quantified using the Qubit dsDNA HS Assay Kit (Invitrogen, US). Ten-fold dilutions were performed on the linearized plasmids, which were used as the standard curve. Besides, we also evaluated the qPCR amplification efficiency using PBFDV-positive fecal samples with 2-fold serial dilution. The amplification efficiencies of the designed assay were 88.6% and 98.6% when the linearized plasmids and PBFDV-positive fecal samples were used as the template DNA, respectively (Supplementary Fig. S1).

## Results

### Variation in viral loads between individuals

Out of 66 fecal samples and 46 back feather samples collected from 17 individuals, 34 (51.5%) and 31 (67.4%) samples were found to be PBFDV-positive using qPCR (Fig. 1). Among 17 individuals, only 1 individual (i.e., RAA21) were found to be PBFDV-negative in both fecal and feather samples. Six individuals were found to be either PBFDV-positive in fecal samples but not feather samples (RAA15-16, and 18), or positive in feather samples but not in fecal samples (RAA22-24).

**Figure 1.**
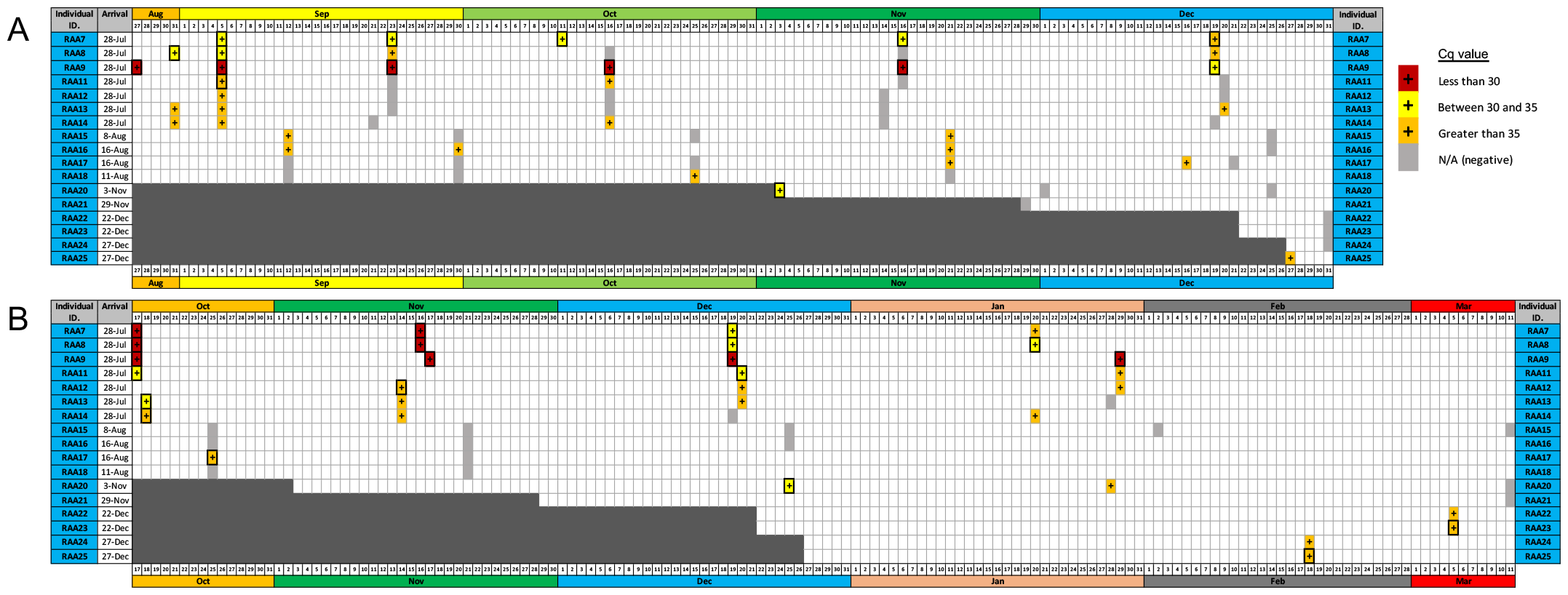
Summary of the PBFDV detection results of 17 *Agapornis roseicollis* individuals. **A**. Fecal samples. **B**. Back feather samples. All samples were tested in duplicate using qPCR. The coloured cells indicated the sample collection date with different colour representing the Cq value of the qPCR result. Only samples that were positive in all technical replicates are marked with a thick box border. “Arrival” refers to the date that the bird started to be housed in an individually ventilated cage.

For individuals with temporal sampling (12 individuals), three individuals (RAA7-9) were consistently PBFDV-positive with a relatively low Cq values in both fecal and feather samples, indicating a large amount of viral DNA copies (Fig. 1). These three individuals were considered to contain a high viral load of PBFDV.

Other individuals with PBFDV-positive samples did not show a consistent result throughout the sampling period and between fecal and feather samples. The positive results for these individuals were occasionally detected with high Cq values (>35 cycles; equivalent to ∼1 copy of PBFDV DNA), indicating low viral load. The chance of false positive for these individuals were low because multiple positives results were detected independently. All negative controls in the qPCR assays showed no PBFDV amplification. These individuals were therefore considered to have a low viral load of PBFDV. Yet, despite the differences in the viral load, all birds did not show any observable PBFD symptoms and behaved normally throughout the sampling period.

### Temporal viral load changes within an individual

Focusing on the individuals with a high viral load (RAA7-9), we further demonstrated that the viral load in both the fecal and feather samples dropped continuously within 200 days after the birds arrived at the facility (Fig. 2 and 3). The trend was similar among all sample types (Fig. 2 and 3). The averaged reduction in viral load was similar between fecal and feather samples (-82.32% and -96.58%, respectively). Notably, the fecal viral load in one individual (RAA8) was barely detectable after ∼3 months (Fig. 3).

**Figure 2.**
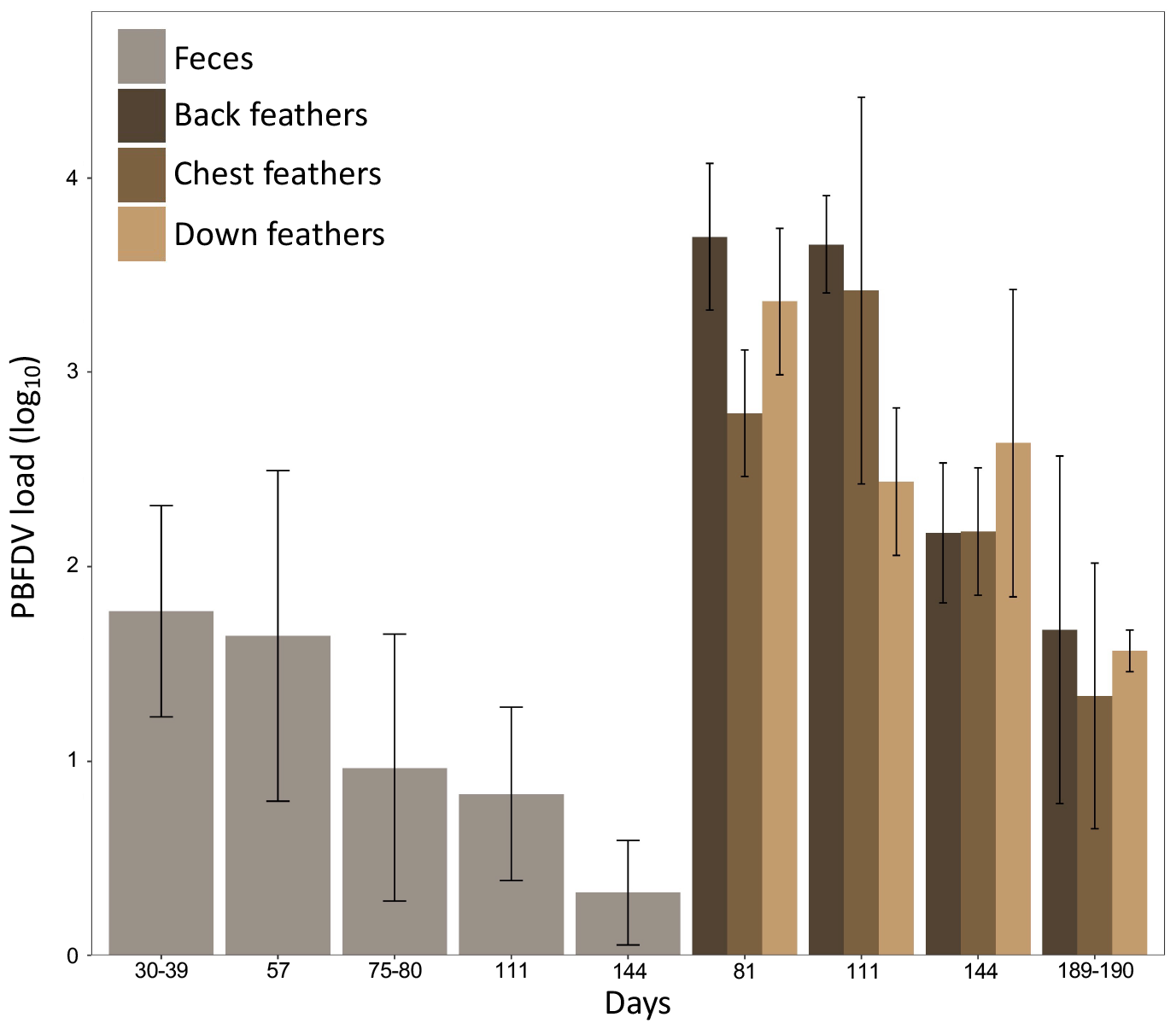
PBFDV viral load of faecal and feather samples in the three *Agapornis roseicollis* individuals that had high viral load at the beginning of the experiment. These three individuals were “RAA7”, “RAA8”, and “RAA9” in Figure 1. The day in the x-axis refers to the day of sample collection after the birds were housed in individually ventilated cages.

**Figure 3.**
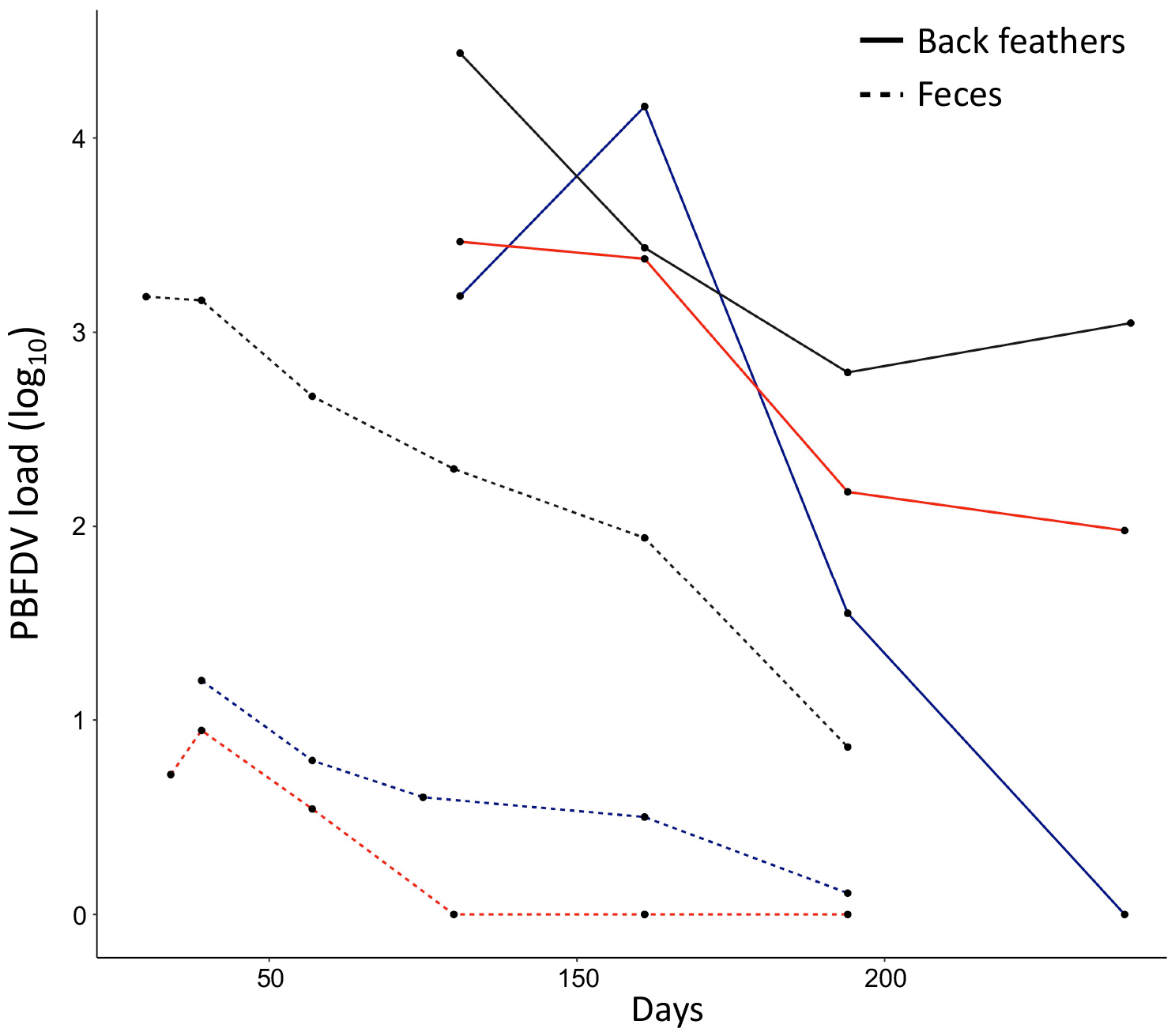
Temporal change of PBFDV viral load in fecal (solid line) and back feather (dashed line) samples in three *Agapornis roseicollis* individuals that had high viral load at the beginning of the experiment. These three individuals were “RAA7” (blue), “RAA8” (red), and “RAA9” (grey) in Figure 1. The day in the x-axis refers to the day of sample collection after the birds were housed in individually ventilated cages.

### Feathers had a higher PBFDV load than feces

For the individuals with a high viral load (RAA7-9), feather viral load (mean = 10^2.73^ copies) were 17.8-fold higher than fecal vial load (mean = 10^1.48^ copies; Fig. 2). At each time point, the viral load in feather samples were higher than that in fecal samples for each individual.

## Discussion

Most individuals in this study were PBFDV-positive but they did not exhibit any observable PBFD symptoms and behaved normally. Notably, despite of high PBFDV prevalence, only three individuals had a relatively high viral load, while most of the other individuals had very low level of viral load. Among the individuals with high viral load, the viral load decreased drastically from a high level to only a few copies detected within 100 days in both feather and fecal samples. In one individual with high viral load at the beginning of the experiment, the viral loads in its fecal samples became barely detectable after ∼3 months.

### The difference in viral load between feather and fecal samples

Feather samples from all the three body regions had a higher PBFDV load than feces. It took longer for the viral load to drop to a relatively low level in feathers than feces. One possible explanation for this finding is that the virus and viral DNA in the feathers represent the historical infection [15]. The virus and viral DNA could persist for a longer period of time in feathers than in the digestive system. Previous studies have shown that viral inclusion bodies in feathers are common during PBFDV infection and feather follicle is one of the major virus replication sites [23]. Virus could accumulate and stay in the feather for a long period, and may only be removed during cleaning (e.g. preening and bathing) or molting. Viral load therefore remains at higher level in feathers than feces, and decreases slower in feathers even after the peak infection phase compared with the faster dropping of viral load in feces (i.e. a proxy of viral load in the gut). Similar findings were observed in budgerigars (*Melopsittacus undulatus*) and crimson rosella [7, *Platycercus elegans*; 15]. Accordingly, it could imply that the viral load in feather samples might not be indicative of the infection status of the bird at the time of sampling. On the other hand, the bursa of Fabricius has been suggested as one of the main sites of PBFDV replication [28], and there are sophisticated immunological interactions between immune cells and pathogens in the gut for the host to clear the invading virus [52]. Viral load in fecal samples therefore could better reflect the current PBFDV infection status than that in feather samples. Given that PBFDV is highly persistent in the environment, the potential for the virus to remain in feathers for an extended period of time can pose a lasting infection risk, even if the host has recovered or suppressed the virus. The potential for recovered or tolerant birds, which show no signs of disease, to carry the virus and shed it into the environment or infect other birds has significant implications for conservation and disease management.

### Rosy-faced lovebirds appears to have a high tolerance to PBFDV

Variations in the susceptibility to PBFDV infection between different species has been observed [20]. Eclectus Parrots (*Ecletus roratus*) and Gang Gang Cockatoos (*Callocephalon fimbriatum*) suffer severe consequences of PBFDV infection, while Cockatiels (*Nymphicus hollandicus*) were said to be less susceptible to the infection [38, 43, 46]. Some lovebird species are highly susceptible with some advanced clinical signs [20, 51]. 100% mortality was reported in captive flocks of the Black-cheeked lovebird (*A. nigrigenis*) and Lilian’s lovebird (*A. lilianae*) [20]. In contrast, our results suggest that *A. roseicollis* is tolerant to PBFDV infection as evidenced by the lack of signs of disease, low viral loads in most PBFDV-positive birds, and fast decrease in viral load in individuals that originally had a high viral load within a short period of time. *A. roseicollis* and *A. fischeri* that were in close contact with diseased birds were observed to show no signs of disease [20]. Other studies also reported recovery of lovebirds from PBFDV-associated feather abnormality, but it is uncertain whether those individuals developed chronic form of the disease or became carriers [31, 34]. Although no PBFDV was found in a wild population of *A. lilianae* in Malawi, South Africa, the virus contact history of the population was unknown [29]. These findings, taken together, suggest that some species might have developed resistance or tolerance to PBFDV infection [26].

We showed that the PBFDV load can drop from a high level to an undetectable level in the feces of rosy-faced lovebirds within a few months, and most of the PBFDV-positive birds shed very little virus into the environment. The birds in this study were infected by PBFDV before entering our housing facility, and the exact timing of infection was not known. Infection experiments could be conducted in the future to gain a deeper understanding of the dynamics of host-virus infections. The discovery of higher viral loads in feathers, which can persist for longer periods compared to feces, has provided crucial insights into the potential risk of PBFDV infection for conservation and disease management purposes. The knowledge can help inform strategies to mitigate the spread of the virus and protect parrot populations.

## Acknowledgements

We thank the CCMR staff, Ellen Sai Nam Lo, Tat Sing Ngai, and Mei Ying Wu for animal care and husbandry. We thanks Jennifer Le Lin Go for comments on the manuscript.

## Statements and Declarations

### Data Availability

The datasets generated during and/or analyzed during the current study are available from the corresponding author on reasonable request.

### Author Contributions

D.K.L. and S.Y.W.S. designed the research. D.K.L. and E.S.K.P. conducted the research. S.Y.W.S. supervised the project. S.Y.W.S. obtained funding for the project. D.K.L. wrote the first draft of the paper. All authors contributed to revised versions.

### Ethics approval

This study was approved the Animal Research Ethics Committee (4751-18) of the University of Hong Kong and the Department of Health [(18-1206) in DH/SHS/8/2/3 Pt. 23] of the HKSAR Government.

